# Broadly-neutralizing antibodies that bind to the influenza hemagglutinin stalk domain enhance the effectiveness of neuraminidase inhibitors via Fc-mediated effector functions

**DOI:** 10.1101/2021.11.17.468248

**Authors:** Ali Zhang, Hanu Chaudhari, Yonathan Agung, Michael R. D’Agostino, Jann C. Ang, Matthew S. Miller

## Abstract

The conserved hemagglutinin stalk domain is an attractive target for broadly effective antibody-based therapeutics and next generation universal influenza vaccines. Protection provided by hemagglutinin stalk binding antibodies is principally mediated through activation of immune effector cells. Titers of stalk-binding antibodies are highly variable on an individual level, and tend to increase with age as a result of increasing exposures to influenza virus. In our study, we show that stalk-binding antibodies cooperate with neuraminidase inhibitors to protect against influenza virus infection in an Fc-dependent manner. These data suggest that the effectiveness of neuraminidase inhibitors is likely influenced by an individual’s titers of stalk-binding antibodies, and that neuraminidase inhibitors may enhance the effectiveness of future stalk-binding monoclonal antibody-based treatments.

## Introduction

Influenza causes 3-5 million serious illnesses and 290,000 – 645,000 deaths every year worldwide (Iuliano et al., 2018). Seasonal influenza vaccination induces production of mostly neutralizing antibodies directed against the hemagglutinin (HA) head domain and is the most effective way to prevent infection. The HA head domain undergoes continuous antigenic drift, which necessitates annual reformulation of seasonal vaccines to narrow strain-specific protection. Novel vaccine strategies designed to elicit immune responses against more conserved protein domains, such as the HA stalk domain by broadly neutralizing antibodies (bNAbs), may achieve more broad and durable protection against influenza.

The HA stalk domain is more conserved between influenza virus strains compared to the HA head domain due to functional constraints and minimal immune pressure (Kirkpatrick et al., 2018; Wu and Wilson, 2017). Protection against influenza by HA stalk-binding antibodies is mediated primarily by Fc-dependent activation of immune effector cells (DiLillo et al., 2016, 2014; He et al., 2016). Optimal Fc activation by these antibodies requires two points of contact: the antibody Fab-HA stalk interaction, and the sialic acid-HA head interaction (Leon et al., 2016) (Figure 1A). Neuraminidase (NA), the other major influenza virus surface glycoprotein, cleaves sialic acids from the HA head domain during viral budding. Antibody dependent cellular cytotoxicity (ADCC) is a process through which infected cells are eliminated by Fc receptor (FcR)-bearing immune cells after binding of antibodies to surface-bound antigen. Antibodies that bind to NA can induce modest levels of ADCC on their own and can also cooperate with stalk-binding antibodies to enhance ADCC induction (He et al., 2016). HA-stalk binding antibodies can, in turn, partially inhibit neuraminidase activity, contributing to their potency of virus neutralization (Kosik et al., 2019). Since cleavage of the sialic acid-HA head interaction by neuraminidase may destabilize one of the two points of contact required for optimal stalk antibody-mediated activation of immune effector cells, we reasoned that chemical inhibition of NA activity may also potentiate ADCC induction by HA stalk-binding antibodies.

**Figure 1.**
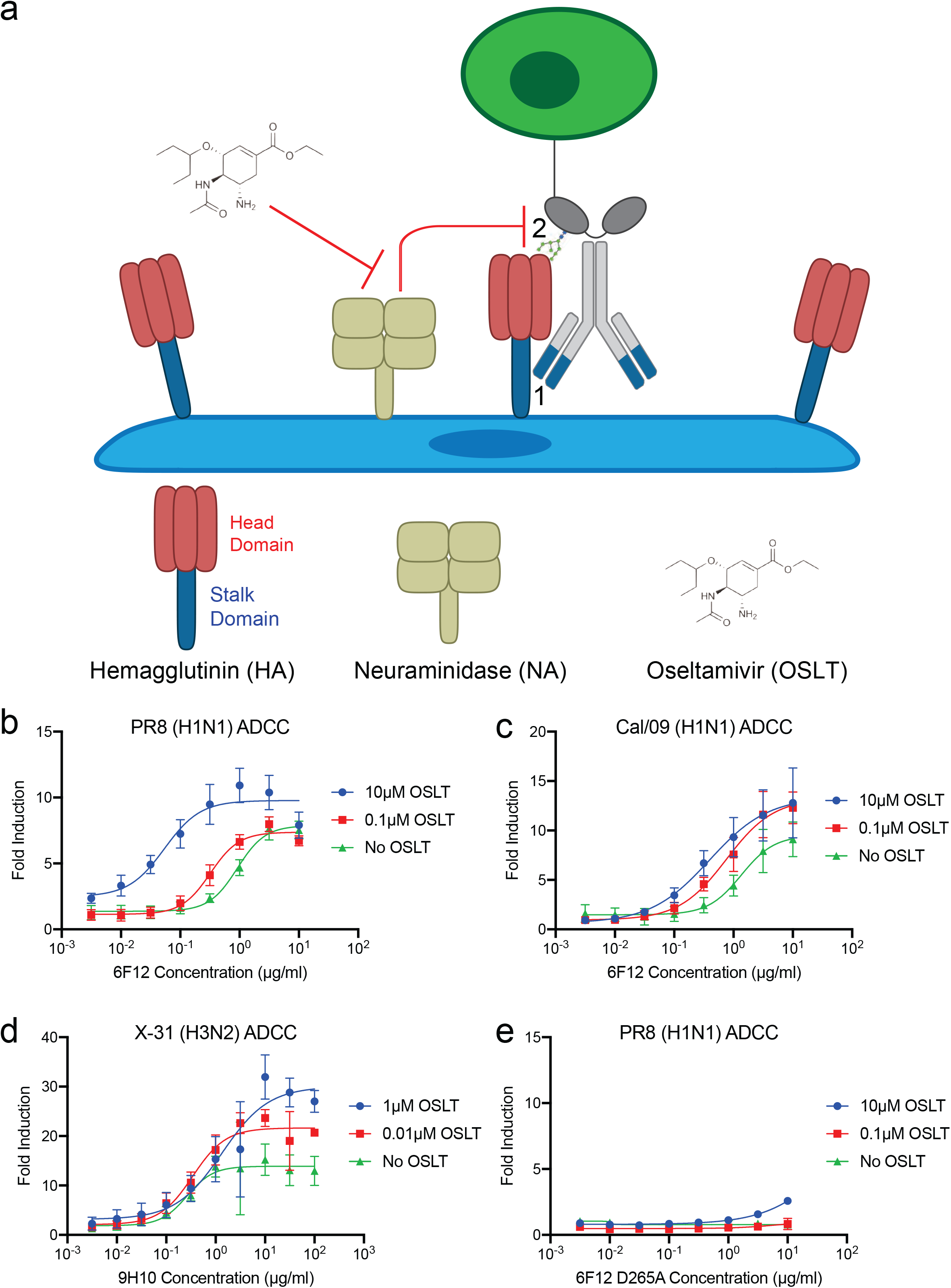
Oseltamivir potentiates ADCC induction by monoclonal stalk-binding broadly-neutralizing antibodies in a dose-dependent manner. (A) Diagram of bNAb facilitating the interaction between immune effector cell and infected cell via two points of contact. The immune effector cell with an Fc receptor is depicted in green. The stalk-binding antibody is shown in grey and blue. The stalk-binding antibody interacts with the HA stalk domain via its Fab portion (1) and binds to the Fc receptor of the effector via its Fc portion. The HA head domain interacts with sialic acid residues on the effector cell (2). NA enzymatically cleaves sialic acids from the HA head domain, abrogating the second point of contact. NA inhibitors, such as oseltamivir, restore the second point of contact by preventing the enzymatic cleavage of HA and sialic acid. (B-E) In vitro ADCC assays were completed using A549 cells infected with PR8 (H1N1), Cal/09 (H1N1) or X-31 (H3N2) at an MOI of 5. Fold induction depicts activation above background (infected cells without antibody). Concentrations of oseltamivir carboxylate are denoted in the legend. 6F12 (Pan H1 stalk-binding antibody) was used to target PR8 and Cal/09 infected cells, while 9H10 (Group 2 HA stalk-binding antibody) was used to target X-31 infected cells. Fold induction data is shown as mean ± SD with biological triplicates.

Here, we show that chemical inhibition of neuraminidase increases stalk antibody Fc-mediated activation of immune effector cells in a dose dependent manner. HA stalk antibodies potentiated the efficacy of the NA inhibitor oseltamivir in both prophylactic and therapeutic contexts. This effect is preserved in the contexts of both monoclonal stalk antibody and polyclonal sera. Cooperativity between neuraminidase inhibition and HA stalk antibodies was dependent on Fc-FcR interaction. Together, these data suggest that the efficacy of oseltamivir treatment may be dependent on HA stalk antibody titers, and that monoclonal HA stalk antibodies and neuraminidase inhibitors may represent an effective combination therapy in the future.

## Results

### Oseltamivir cooperates with stalk-binding antibodies to enhance antibody-dependent cell cytotoxicity of cells infected with group 1 and group 2 influenza A viruses

We first set out to determine the role of neuraminidase on HA stalk-binding antibody mediated activation of immune effector cells. We first quantified the NA activity of three representative strains of influenza virus, A/Puerto Rico/8/1934 H1N1 (PR8), A/California/07/2009 H1N1 (Cal/09), and X-31 H3N2 (Figures S1A-S1C) (Kilbourne, 1969). We then tested the potency of oseltamivir in inhibiting the NA activity of these strains (Figures S1D-F), and in reducing viral replication (Figures S1G-S1I). Having established these dose-response relationships with oseltamivir, we performed antibody dependent cell cytotoxicity (ADCC) assays using modified Jurkat effector cells expressing murine FcγRIV. This is an extremely sensitive, highly quantitative assay that we have used extensively in the past and has been validated to correlate closely with the activation of primary NK cells (Chromikova et al., 2020; He et al., 2016). A549 target cells were infected with PR8, Cal/09, or X-31 at an MOI of 5. The infected cells were then incubated with oseltamivir and murine monoclonal stalk-binding broadly neutralizing antibodies (bNAbs) 6F12 for cells infected with PR8 and Cal/09 (which express group 1 HA), and 9H10 for cells infected with X-31 (which expresses group 2 HA). Infection and binding specificity of antibodies was verified by immunofluorescence microscopy (Figure S2). Addition of oseltamivir resulted in a dose-dependent enhancement of ADCC induction in PR8, Cal/09 and X-31 infected cells (Figures 1B-D). Enhancement of bNAb-mediated ADCC by oseltamivir required Fc-FcR interaction, as 6F12 with the D265A mutation, which abrogates FcR binding, failed to induce ADCC of PR8 infected cells, even at the highest oseltamivir concentration used (Figure 1E). Together, these results demonstrate that NA inhibition by oseltamivir enhances ADCC induction by bNAbs that bind to the HA stalk domain, demonstrating cooperative activation.

### The efficacy of oseltamivir in preventing and treating influenza virus is potentiated by bNAbs

To determine whether bNAbs could enhance protection mediated by oseltamivir *in vivo*, we conducted influenza challenge experiments in a murine model system. To test efficacy in a prophylaxis setting, six groups of 6-8 week old BALB/c mice (n = 5 mice/group) were administered 1mg/kg 6F12, 6F12 D265A, or IgG isotype control intraperitoneally and 1mg/kg oseltamivir or PBS by oral gavage. The mice were then infected with 5 × LD_50_ of PR8 (500 plaque-forming units (PFU)) intranasally two hours later. Mice continued to receive either oseltamivir or PBS twice daily by oral gavage for 5 days and were monitored for 14 days, with the humane endpoint defined as 80 % of initial body weight (Figures 2A-C). Mice that received negative control treatments all reached endpoint by day 7 post-infection. Of the mice that did not receive oseltamivir, treatment with 1 mg/kg 6F12 resulted in similar disease progression compared to the IgG isotype control. Similarly, mice that received oseltamivir alone experienced significant weight loss. In contrast, mice that received both oseltamivir and 6F12 lost minimal weight throughout the experimental period, with none reaching endpoint. Mice that received both oseltamivir and 6F12 D265A displayed similar disease progression compared to mice that received oseltamivir alone, demonstrating that Fc-FcR interactions are crucial for the enhanced protection offered by co-administration of 6F12 and oseltamivir.

**Figure 2.**
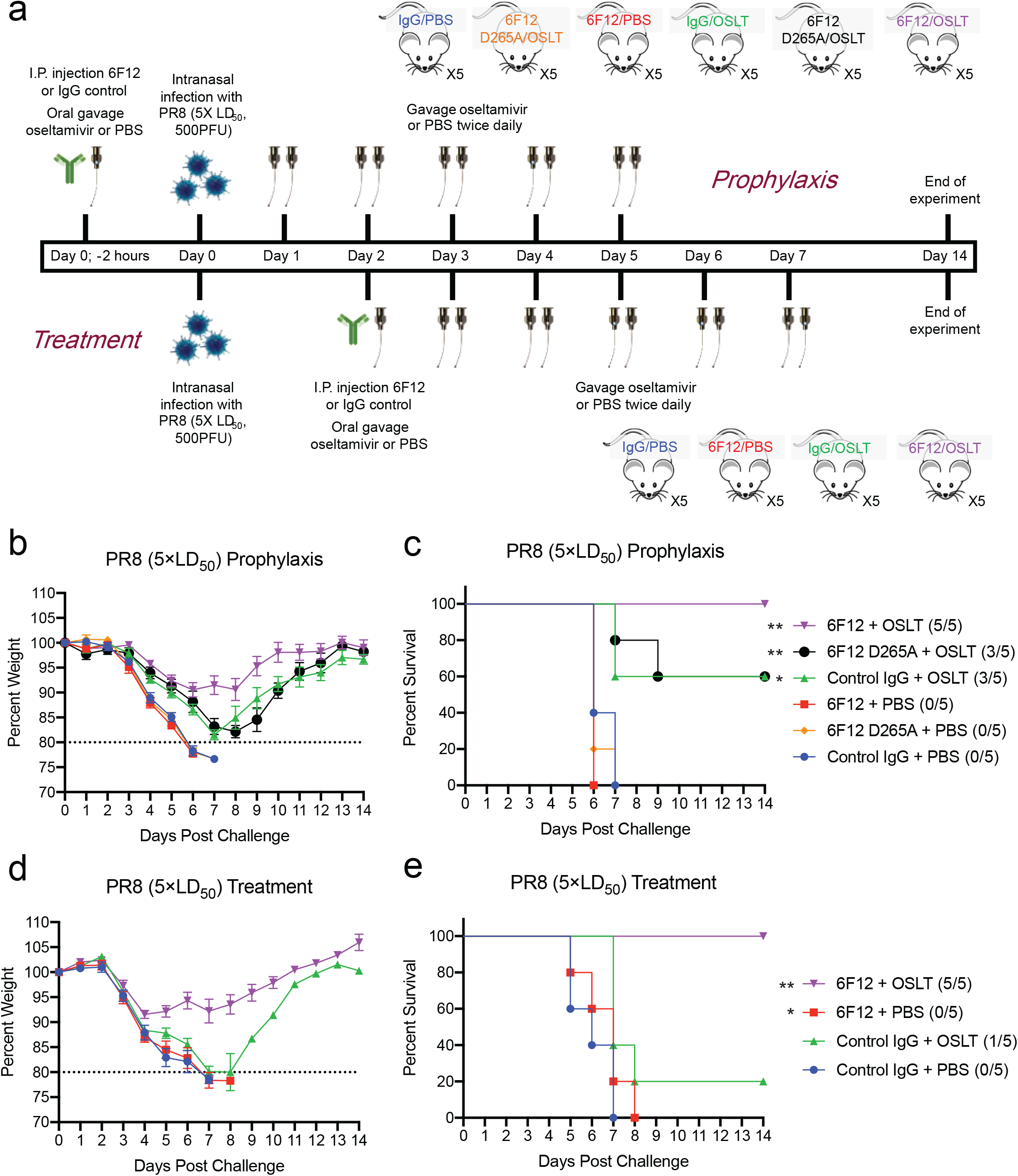
Oseltamivir administered in combination with bNAbs is superior at preventing and treating influenza clinical signs compared to either therapeutic alone. (A) 6-8 week old female BALB/c mice were infected intranasally with 500 PFU of PR8 (5 × LD_50_). The mice were also administered intraperitoneally with 6F12 or PBS and an oral gavage of oseltamivir or PBS. A dose of either 1mg/kg 6F12 and/or 1mg/kg oseltamivir (prophylaxis) or 10mg/kg 6F12 and/or 10mg/kg oseltamivir (treatment) were used. The first round of therapeutics was given either 2 hours before infection (prophylaxis) or 48 hours after infection (treatment). Mice were then given oseltamivir or PBS by oral gavage twice daily for 5 additional days following the first round of therapeutics. Weight change was monitored daily, and the animals were sacrificed when they reached 80% of initial weight. (B-C) Weight loss and Kaplan-Meier survival curves of the mice treated prior to infection (prophylaxis group). Weight loss is shown as percent of initial weight with mean ± SEM, n=5/group. Statistical comparisons are shown against the control IgG + PBS group; *p<0.05, **p<0.01. Numbers in brackets denote number of surviving mice and total number of mice per group. (D-E) Weight loss and Kaplan-Meier survival curve of the mice treated after infection (treatment group). Weight loss is shown as percent of initial weight with mean ± SEM, n=5/group. Statistical comparisons are against the control IgG + PBS group; *p<0.05, **p<0.01. Numbers in brackets denote number of surviving mice and total number of mice per group.

To determine if bNAbs could also enhance the activity of oseltamivir when treating an established influenza infection, we infected four groups of 6-8 week old BALB/c mice (n = 5 mice/group) with 5 × LD_50_ of PR8 (500 PFU) and treated with 10mg/kg 6F12 or 10mg/kg oseltamivir starting 2 days post-infection (Figures 2D-E). Antibody or oseltamivir alone were insufficient to protect mice from severe disease, whereas the combination of 6F12 and oseltamivir yielded complete protection. These results demonstrate a substantial improvement in protection against influenza morbidity and mortality when oseltamivir and stalk-binding antibodies are administered together in the context of both prophylaxis and treatment. The cooperativity of oseltamivir and stalk-binding antibodies was not due to their independent activities (i.e., direct neuraminidase inhibition and virus neutralization), but was instead dependent upon Fc-FcR interactions mediated by the stalk-binding antibodies.

### Titers of bNAbs in human serum predict effectiveness of oseltamivir treatment

The previous results demonstrate that oseltamivir cooperates with monoclonal stalk-binding antibodies to protect against clinical signs of influenza virus infections by enhancing antibody Fc-mediated immune effector cell activation. We next set out to determine if titers of bNAbs in human serum could predict the effectiveness of oseltamivir treatment. First, we conducted ADCC assays using serum from 4 adult donors who had received the seasonal influenza vaccine 14 days prior to serum collection. Titers of bNAbs in these samples were first quantified by ELISA using a chimeric cH6/1 hemagglutinin protein, that contains the head domain from A/Mallard/Sweden/81/02 H6N1 and the stalk domain from A/Puerto Rico/8/1934 H1N1 (Figures 3A and 3B). The cH6/1 chimera has been used extensively in the past to quantify antibodies that specifically bind to the group 1 stalk domain, as humans are rarely exposed to H6 influenza viruses and we have previously shown that human sera lacks hemagglutination inhibition (HAI) activity against the H6 head domain (Miller et al., 2013b; Nachbagauer et al., 2014; Pica et al., 2012; Sangster et al., 2013). Consistent with the observations made using monoclonal bNAbs, ADCC induction by polyclonal bNAbs from human donors was enhanced by oseltamivir (Figures 3C–3F).

**Figure 3.**
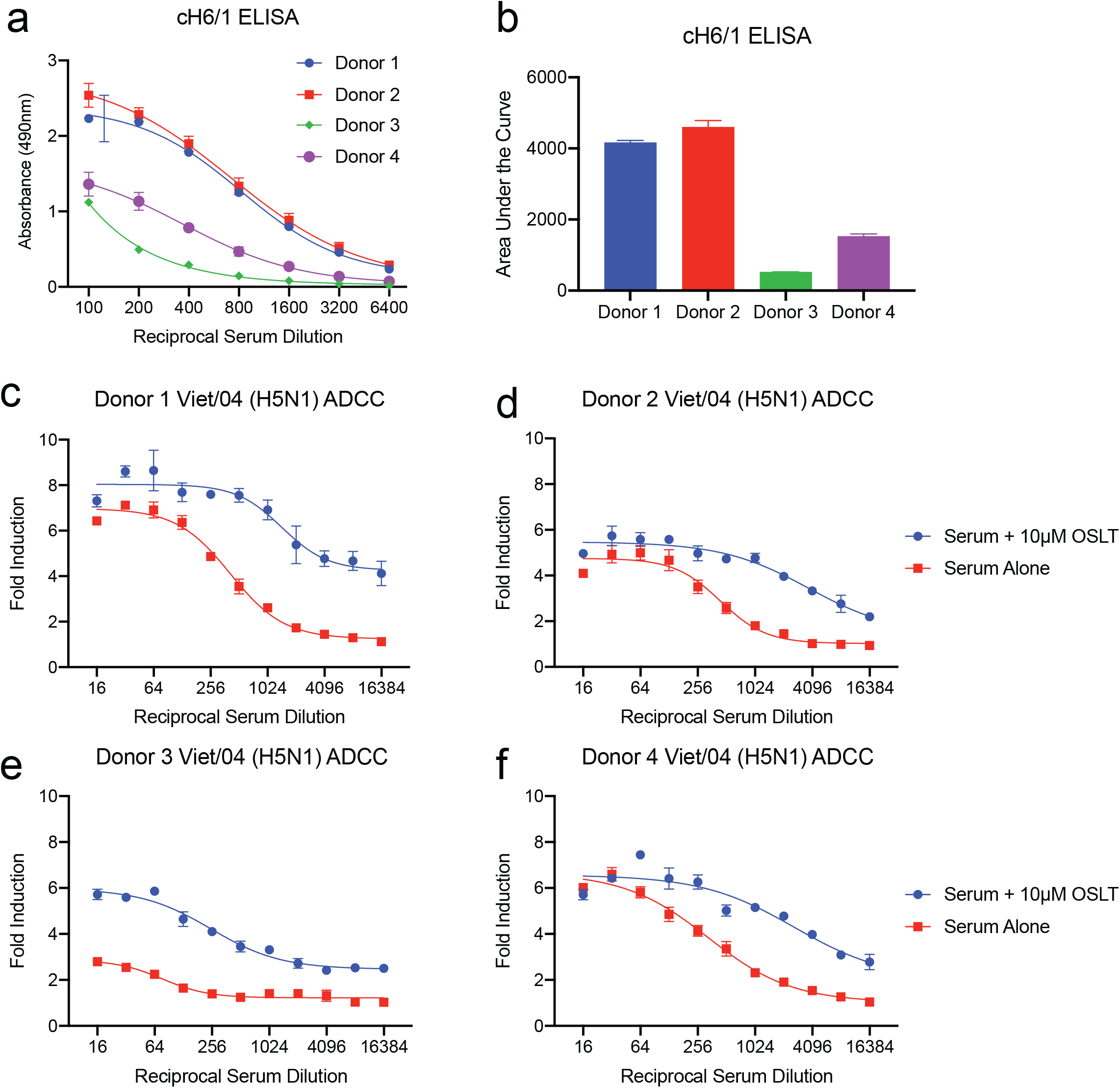
Oseltamivir increases the potency of ADCC induction by polyclonal stalk-binding antibodies. (A-B) ELISAs were performed using recombinant cH6/1 HA to measure the titer of H1 stalk-binding antibodies in the serum samples. Absorbance data is shown as mean ± SD. The area under the curve is also shown as mean ± SEM. (C-F) In vitro ADCC assays were completed using A549 cells infected with Viet/04 (H5N1) at an MOI of 5. Fold induction denotes activation above infected cells without antibody. Serum was obtained from 4 healthy donors who had been previously vaccinated with seasonal influenza virus vaccine. Fold induction data is shown as mean ± SD.

We next set out to assess the effectiveness of human sera in combination with oseltamivir in protecting against a representative “pandemic-like” virus. We then chose to use A/Vietnam/1203/2004 H5N1 HAlo (Viet/04), which is a an H5N1 virus that is highly pathogenic in mice, but has had the polybasic cleavage site of the H5 protein removed to enhance safety (Steel et al., 2009). ELISAs were first performed on the serum from two additional healthy adult donors using the chimeric cH6/1 hemagglutinin protein to quantify bNAb titers (Figures S3A and S3B)., To ensure that the serum samples used in these assays did not contain HAI+ antibodies to the head domain of Viet/04 H5, we conducted a HAI assay (Figure S3C). No HA inhibition was observed, indicating that both donors were naïve to this virus. In ADCC assays, the donor with higher titers of HA stalk-binding antibodies based on ELISA data (Figure S3A) also elicited higher activation of reporter cells, consistent with expectations (Figure S3D).

Next, six groups of 6-8 week old BALB/c mice were either administered 150ul of serum with low titers of stalk-binding antibodies (Low Serum), 150ul of serum with high titers of stalk-binding antibodies (High Serum), or 150ul of PBS (vehicle). The mice were also given either 0.1mg/kg oseltamivir or PBS by oral gavage. The mice were infected with 5 × LD_50_ of Viet/04 H1N1 (200 PFU) intranasally two hours after passive transfer of serum (Figure 4A). The data is displayed as two sets of graphs for clarity, with shared negative control groups (Figures 4B–4E). Mice that received either serum or oseltamivir alone experienced significant morbidity, with most animals in those groups reaching endpoint. When oseltamivir was administered to mice that had received serum containing low titers of bNAbs (Low Serum), weight loss was similar to that of mice treated with serum or oseltamivir alone; however, mortality in this group improved substantially, with only 1 out of 5 mice reaching endpoint. In contrast, when oseltamivir was administered to mice that received serum containing high titers of bNAbs (High Serum) minimal weight loss was observed, and full protection against mortality was achieved (Figures 4B–4E). Taken together, these results demonstrate that titers of bNAbs found in human serum can significantly impact the efficacy of oseltamivir treatment.

**Figure 4.**
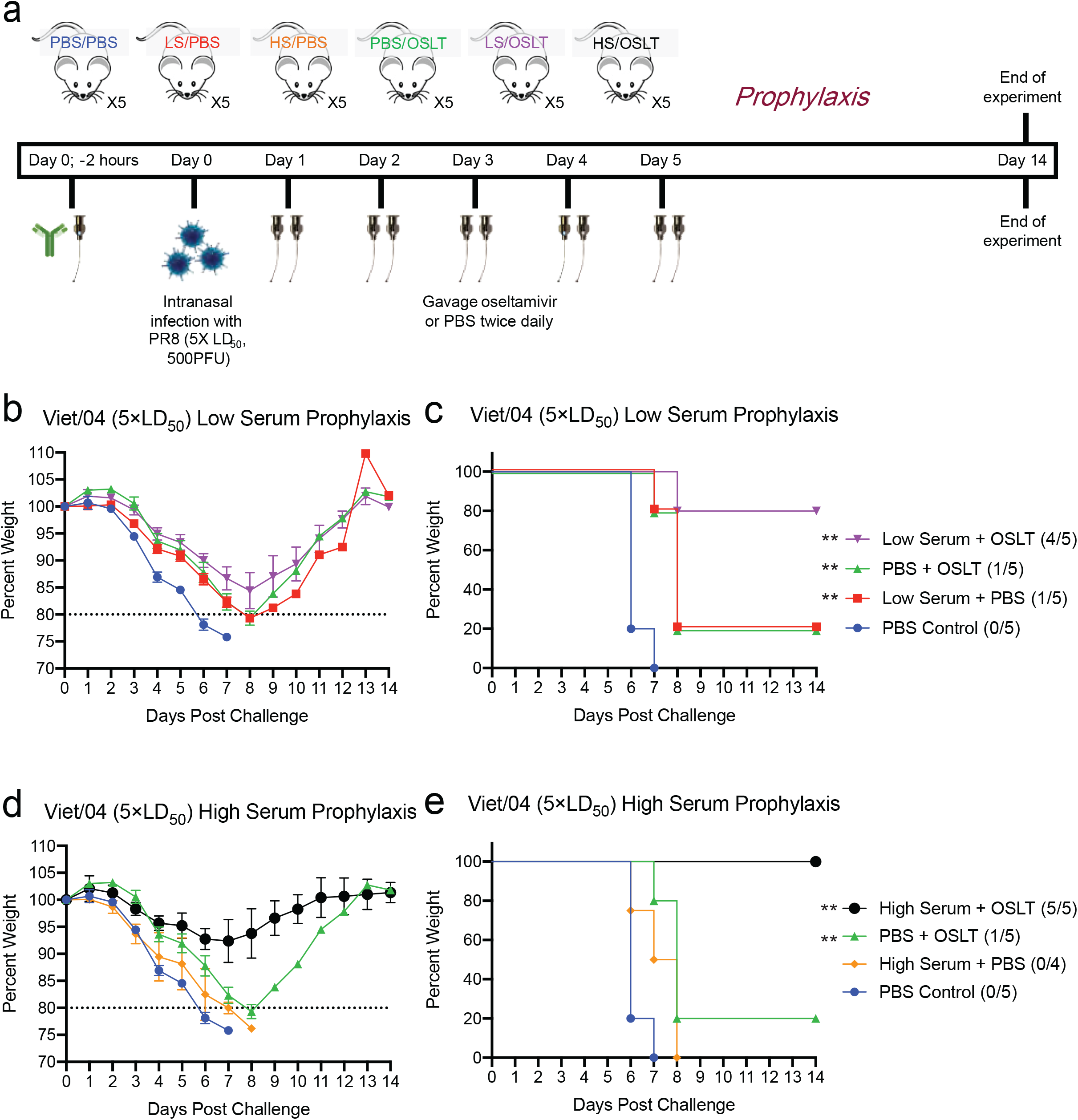
Oseltamivir is more effective at preventing influenza clinical signs in animals with higher HA stalk-binding antibody titers. (A) 6-8 week old female BALB/c mice were administered 150ul of human serum or PBS i.p. and an oral gavage of 0.1mg/kg of oseltamivir or PBS. The serum contains either high titers polyclonal stalk-binding antibodies (High Serum; HS), or low titers of polyclonal stalk-binding antibodies (Low Serum; LS). The mice were then infected intranasally with 200 PFU of Viet/04 (5 × LD_50_) two hours later. Mice were given oseltamivir or PBS by oral gavage twice daily for 5 days. Weight change was monitored daily, and the animals were sacrificed when they reached 80% of initial weight. (B-C) Weight loss and Kaplan-Meier survival curve of the Low Serum group. Weight loss is shown as percent of initial weight with mean ± SEM, n=5/group. Statistical comparisons are against the PBS Control group; *p<0.05, **p<0.01. PBS+OSLT and PBS Control groups are shared between graphs depicting High Serum and Low Serum experiments for clarity. Numbers in brackets denote number of surviving mice and total number of mice per group. (D-E) Weight loss and Kaplan-Meier survival curve of the High Serum group. Weight loss graph is shown as percent of initial weight as mean ± SEM, n=5/group unless otherwise indicated. Statistical comparisons are against the PBS Control group; *p<0.05, **p<0.01. PBS+OSLT and PBS Control groups are shared between graphs depicting High Serum and Low Serum experiments for clarity. Numbers in brackets denote number of surviving mice and total number of mice per group.

## Discussion

Neuraminidase inhibitors are a major class of antivirals used for treatment and prevention of influenza virus. While NA inhibitors tend to be highly effective in preventing influenza when administered prophylactically, effectiveness is much more modest in most therapeutic contexts (Doll et al., 2017). Maximum benefit is achieved therapeutically when NA inhibitors are administered early after symptom onset (Fry et al., 2014; Hiba et al., 2011; Rodríguez et al., 2011). However, there is considerable heterogeneity in the effectiveness of these drugs at the individual level, and the host factors that determine outcome remain elusive (Jefferson et al., 2014). Furthermore, neuraminidase inhibitor-resistant viruses circulate naturally, and therefore strategies that mitigate the selection of these resistant variants are critical to prolonging the utility of this class of drug (Lampejo, 2020).

The hemagglutinin stalk domain is an attractive target for antibody-based therapeutics and universal influenza vaccines due to its high degree of conservation amongst diverse influenza viruses. The main mechanism by which these antibodies protect against influenza is through Fc-dependent effector cell functions, such as ADCC, NETosis, phagocytosis, and secretion of cytokines (DiLillo et al., 2016, 2014; He et al., 2017; Stacey et al., 2021). Elegant studies by the Yewdell laboratory previously demonstrated that NA inhibition by anti-HA stalk antibodies contributes to viral neutralization and induction of FcγR-mediated activation of innate immune cells (Kosik et al., 2019). We have also previously shown that antibodies with NAI activity can cooperate with bNAbs that bind to the HA stalk to potentiate ADCC (He et al., 2016). Our current study demonstrates that the efficacy of NA inhibitors is profoundly influenced by pre-existing titers of bNAbs, and that protection from influenza *in vivo* is mediated by potentiation of Fc-dependent effector functions of immune cells mediated by NA inhibition.

We used the well-characterized monoclonal murine antibodies 6F12 and 9H10, which bind to H1 and group 2 HA respectively, in our ADCC assays (Tan et al., 2014, 2012). Our ADCC assay uses engineered Jurkat effector cells, which express murine FcγRIV and firefly luciferase driven by the nuclear factor of activated T cells (NFAT) response element. ADCC induction in these assays correlate well with classical CD107a NK cell degranulation assays (Chromikova et al., 2020). We found that oseltamivir potentiated ADCC induced by bNAbs in a dose-dependent manner. Intact Fc-FcR interactions were absolutely necessary for this potentiation, as introduction of a D265A mutation in the CH2 domain of 6F12, which abolishes the Fc-FcR interaction, negated effector cell induction (Baudino et al., 2008; Nimmerjahn et al., 2005; Temming et al., 2020). The addition of oseltamivir had similar effects in improving ADCC mediated by serum from healthy donors. Effector cell activation is strikingly enhanced even in samples with low titers of bNAbs. Antibodies targeting the less immunogenic NA, M2, and NP have also been shown to induce effector cell activation (DiLillo et al., 2016; Jegaskanda et al., 2017; Lee et al., 2014; Stadlbauer et al., 2019). Our experiments here show that chemical inhibition of neuraminidase can further enhance induction of ADCC even in a complex, polyclonal context.

Consistent with previous studies, we found that co-administration of oseltamivir with a sub-protective dose of monoclonal HA stalk-binding antibodies improve protection against lethal influenza challenge in mouse models (Nakamura et al., 2013; Paules et al., 2017). However, our work clearly demonstrates that this enhancement is dependent upon Fc-FcR interactions, and is not due to the combined, independent activities of oseltamivir and bNAbs. To test whether titers of bNAbs in human serum might impact the effectiveness of oseltamivir treatment, we passively transferred mice with human sera containing either “low” or “high” titers of bNAbs. The effectiveness of oseltamivir treatment was proportional to the titers of bNAbs transferred to the mice. Together, these data strongly suggest that and individual’s pre-existing titers of bNAbs may have a profound impact on the effectiveness of oseltamivir treatment.

Our data provide new insights that help to explain the heterogeneity in effectiveness of NA inhibitors at the individual level. In the future, human clinical trials that directly assess whether pre-existing titers of bNAbs predict the effectiveness of NA inhibitor treatment could have important clinical implications that inform selection of the most appropriate antiviral treatments for individuals with influenza – especially in light of the recent approval of baloxivir marboxil, which targets the cap-snatching endonuclease activity of the viral polymerase complex (Noshi et al., 2018). These drugs have similar clinical effectiveness and thus, an evidence-based framework to guide which drug may be most effective for individual patients is sorely needed. While rapid screening of bNAb titers may be possible in the future, it is also known that titers of these antibodies tend to increase with age (Miller et al., 2013a; Nachbagauer et al., 2016). Our study also suggests that the effectiveness of future bNAb-based monoclonal antibody therapies could be improved by coadministration with NA inhibitors. This class of drugs may be especially important for providing first-line defense in the event of a future influenza virus pandemic, since prediction of which strains are likely to cause pandemics remains a considerable challenge to strain-specific pandemic vaccine design (Miller and Palese, 2014).

## Acknowledgements

This work was supported, in part, by a Canadian Institutes of Health Research operating grant, a Boris Family Foundation grant, and an M.G. DeGroote Institute for Infectious Disease Research seed funding grant (M.S.M). M.S.M was also supported, in part, by a CIHR Early Investigator Award and an Ontario Early Researcher Award. M.S.M. holds a Canada Research Chair in Viral Pandemics. A.Z is supported by a Physician Services Incorporated Research Trainee Fellowship and a CHIR Canada Graduate Scholarships – Doctoral Award. M.G.D. was supported, in part, by an Ontario Graduate Scholarship.

## Author Contributions

Conceptualization, A.Z. and M.S.M.; Methodology, A.Z. and M.S.M.; Formal Analysis, A.Z., H.C., Y.A., J.C.A., and M.S.M; Investigation, A.Z., H.C., Y.A., M.R.D., and J.C.A.; Writing, A.Z. and M.S.M.; Supervision, M.S.M

---

**Figure S1** Oseltamivir inhibits neuraminidase activity and replication of H1 and H3 influenza viruses.

(A-C) Neuraminidase activity of influenza viruses PR8, Cal/09, and X-31 were measured in-vitro using the NA-Star Neuraminidase Kit (ThermoFisher). Data shown as mean ± SD of at least two technical replicates.

(D-F) Oseltamivir susceptibility of the strains were then determined using 1×10^6^ PFU of the three strains using the NA-Star Neuraminidase Kit (ThermoFisher). The IC50 values are displayed to the right of the graphs. Data shown as mean ± SD of three technical replicates.

(G-I) Oseltamivir sensitivity of influenza virus strains was also quantified using plaque assays by infecting A549 cells with the virus at an MOI of 5 before incubating the infected cells with indicated concentrations of oseltamivir for 18 hours. N.D: not detectable. The limit of detection for the plaque assays was 25 PFU/ml, denoted by the horizontal dotted line. Data shown as mean ± SD of three technical replicates.

**Figure S2** Immunostaining of infected A549 cells by bNAbs.

A549 cells infected with PR8 or Cal/09 were stained with 6F12 or 6F12 D265A, and cells infected with X-31 were stained with 9H10. Uninfected A549 cells were stained using 6F12, 6F12 D265A, and 9H10 as negative controls. Scale bars = 400 μm.

**Figure S3** Characterization of polyclonal stalk-binding antibodies in human serum. Human serum was obtained from peripheral blood of two healthy adult donors.

(A-B) ELISAs were performed using chimeric cH6/1 protein to quantify the titers of antibodies that bind to the stalk domain of H1 hemagglutinin. Absorbance data is shown as mean ± SD with technical triplicates. The area under the curve is also shown as mean ± SEM.

(C) Hemagglutinin inhibition assays using were used to test for the presence of antibodies inhibit receptor binding by the head domain of Viet/04 H5. N.D. = Not detectable.

(D) In vitro ADCC assays were completed using A549 cells infected with Viet/04 (H5N1) at an MOI of 5. Fold induction denotes activation above background (infected cells without antibody). Fold induction data is shown as mean ± SD with technical triplicates.

## Materials and Methods

### Cells and Viruses

Human adenocarcinoma alveolar basal epithelial cells (A549) and Madin-Darby Canine Kidney cells (MDCK) originally obtained from the ATCC were grown in “complete DMEM”, containing DMEM supplemented with 10% (vol/vol) heat-inactivated FBS (Gibco), 100U/mL Penicillin-Streptomycin (Gibco), 2mM GlutaMAX Supplement (ThermoFisher), 10mM HEPES. PR8, X-31, and Viet/04 H5N1 HAlo viruses were propagated in specific pathogen free embryonated chicken eggs (Canadian Food Inspection Agency) (Kilbourne, 1969; Steel et al., 2009). Cal/09 H1N1 was propagated in MDCK cells.

### NA-Star Neuraminidase Assays

NA-Star assays were completed according to manufacturer’s protocol (Invitrogen). Briefly, viruses were diluted in NA-Star Assay Buffer in white 96 well opaque flat-bottom plates to a final volume of 25 μl per well. An additional 25 μl per well of oseltamivir carboxylate (Toronto Research Chemicals) diluted in NA-Star Assay Buffer was added in half-log dilutions to each well. Plates were then incubated for 20 minutes at 37 °C after brief shaking. After the incubation, 10 μl of NA-Star Substrate was added to each well and was then incubated for 30 minutes at room temperature after a brief shaking. After the incubation, 60 μl of NA-Star Accelerator was added to each well, and luminescence was quantified using a SpectraMax i3 plate reader (Molecular Devices).

### Virus Quantification by Plaque Assay

MDCK cells were seeded at a density of 1×10^6^ cells per well in 6 well plates. The next day, cells were infected with serial dilutions of virus diluted in 1× minimum essential medium (MEM, Sigma) supplemented with 0.6% BSA (Sigma). After the 1-hour infection at 37°C, the media was replaced with “Flu Media” containing 1x MEM, 0.6 % BSA, 1μg/ml TPCK-treated trypsin, 0.01% DEAE-dextran, and 0.5% agar. The infected cells were then incubated at 37 °C for 2 days. After the incubation, the cells were fixed with 10 % buffered formalin for 30 minutes at room temperature. The agar layer was then removed, and the cells were stained with crystal violet to visualize plaques.

### ADCC Reporter Assay

A549 cells were seeded at a density of 2×10^4^ cells per well in white 96 well opaque flat-bottom plates in complete DMEM. 24 hours after seeding, cells were infected with PR8, Cal/09, X-31, or Viet/04 at an MOI of 5. 16 hours after infection, the media was replaced with 50 μl of assay buffer (RPMI 1640 supplemented with 4% (vol/vol) low IgG FBS) containing oseltamivir carboxylate (Toronto Research Chemicals) and serial dilutions of monoclonal antibodies or serum. After a 30-minute incubation at 37 °C, 25 μl of 7.5×10^4^ Jurkat effector cells expressing either human FcγRIIIa or murine FcγRIV (Promega) resuspended in assay buffer were added to each well. The cells were incubated for an additional 6 hours at 37°C before 75μl of Bio-Glo Luciferase Assay Reagent (Promega) was added to each well. After a 5-minute incubation at room temperature, luminescence in relative light units (RLU) was quantified using a SpectraMax i3 plate reader (Molecular Devices). Fold induction was obtained by dividing the RLU of the wells of interest by the mean of control wells containing infected target cells and Jurkat effector cells with no monoclonal antibodies/serum.

### Immunofluorescent Staining

A549 cells were seeded at a density of 2×10^4^ cells per well in transparent 96 well flat-bottom plates in complete DMEM. 24 hours after seeding, cells were infected with PR8, Cal/09, or X-31 at an MOI of 5. 16 hours after infection, the cells were fixed using 10 % buffered formalin for 30 minutes at room temperature. After 3 washes with PBS, cells were stained with 6F12, 6F12 D265A, or 9H10 at a concentration of 10 μg/ml diluted in PBS for one hour. After 3 washes with PBS, the cells were then stained using 0.5 μg/ml Goat anti-mouse IgG (H+L) secondary antibody Alexa Fluor 488 (Invitrogen) diluted in PBS for one hour. After 3 washes with PBS, the cells were stained for 5 minutes with 1μg/ml Hoechst 33342 (Life Technologies). The cells were washed again with PBS, and images were taken on an EVOS FL Cell Imaging System (ThermoFisher).

### Antibody Purification

Murine 6F12 and 9H10 were obtained from hybridoma cultures as previously described (Tan et al., 2014, 2012). Hybridomas were thawed and expanded in ClonaCell-HY Growth Medium E (Stem Cell Technologies) up to a volume of 300ml. The media was then changed to Hybridoma-SFM (Gibco). The hybridoma cultures were harvested by centrifugation at 3000 × g for 20 minutes when cultures reached maximal cell density. The supernatant was then nutated overnight at 4 °C with 1 ml of Protein G Sepharose 4 Fast Flow slurry (Invitrogen) per 25ml of supernatant. After incubation, the supernatant/Sepharose bead slurry was passed through a 5 ml polypropylene gravity flow column (Qiagen). The column was washed with 1 column volume of PBS before being eluted with 9 ml of Elution Buffer (0.1M Glycine/HCl buffer, pH 2.2) into 1 ml of Neutralization Buffer (2M Tris-HCl, pH 10). The eluate was then concentrated and buffer exchanged to PBS using 30 kDa Amicon Ultra-15 Centrifugal Filter Units (Millipore) according to the manufacturer’s instructions.

The variable heavy and light chain sequences of 6F12 were cloned into the EcoRI/NheI sites of pFUSEss-CHIg-mG2a (D265A, Invivogen) and the EcoRI/BstAPI sites of pFUSE2ss-CLIg-mk (Invivogen) respectively. The Expi293 Expression System was used to produce the 6F12 D265A antibody according to the manufacturer’s protocols (ThermoFisher). A 2:1 molar ratio of light to heavy chain plasmids was used to transfect Expi293F cells. Antibodies were purified from supernatant as described above.

### Mouse Infections

6-8 week old female BALB/c mice (Charles River) were anesthetized with isoflurane and intranasally infected with 40 μl of PR8 (500 PFU, 5 × LD_50_) or Viet/04 (200 PFU, 5 × LD_50_) diluted in PBS. The mice were administered 1 mg/kg or 10 mg/kg 6F12, 150μl of undiluted serum, or 1mg/kg or 10mg/kg Mouse IgG Isotype Control (ThermoFisher) intraperitoneally either 2 hours before infection, or 48 hours post infection as described in the main text. The mice also received 1 mg/kg or 10 mg/kg oseltamivir phosphate (Toronto Research Chemicals) by oral gavage as described in the main text. Mice were monitored daily and were sacrificed if they reached endpoint, defined as 20 % reduction in initial body weight. All animal experiments were approved by the Animal Research Ethics Board at McMaster University.

### ELISA

Purified soluble cH6/1 comprised of the HA head domain of A/Mallard/Sweden/81/02 H6N1 and the stalk domain of PR8 with a C-terminal T4 trimerization domain and a 6-His tag was generated using a baculovirus expression system as previously described (Pica et al., 2012). Clear flat-bottom 96-well Immulon 4 HBX plates (ThermoFisher) were coated with 50 μl of 2 μg/ml of cH6/1 diluted in ELISA coating buffer (50 mM Na_2_CO_3_, 50 mM NaHCO_3_, pH 9.4) overnight at 4 °C. After the incubation, the plates were blocked for 1 hour with 100 μl of blocking buffer (5 % milk powder in PBS with 5% Tween-20 (PBS-T)). Serum serially diluted in blocking buffer was then added to the wells and incubated for 2 hours at room temperature. After the incubation, the plate was washed 3 times with PBS-T. Anti-human IgG (Fab specific) – peroxidase-conjugated antibody (Sigma) diluted 1:5000 in blocking buffer was added to the wells and incubated for 1 hour at room temperature. After 3 additional washes with PBS-T, 50μl of reconstituted SIGMAFAST OPD (Sigma) was added to each well. The reaction was stopped 10 minutes later by adding 50 μl of 3M HCl to each well, and the absorbance at 490nm was read using a SpectraMax i3 plate reader (Molecular Devices).

### Hemagglutinin Inhibition Assay

Chicken red blood cells diluted 1:1 in Alsever’s solution (Canadian Food Inspection Agency) were diluted 1:10 in PBS and centrifuged at 300 × g for 5 minutes. The supernatant was removed, and 125 μl of the red blood cell (RBC) pellet was added to 25 ml of PBS to make 0.5 % chicken RBC. Serum samples were inactivated by trypsin-heat-periodate treatment by first adding 10 μl of 8 mg/ml trypsin to 20 μl of serum before heating at 56 °C for 30 minutes. After cooling the serum to room temperature, 60 μl of 0.011M KIO_4_ was added followed by a 15-minute incubation at room temperature. The periodate treatment was inactivated by adding 60 μl of 1% glycerol (v/v) in PBS followed by a 15-minute incubation at room temperature. Lastly, 50 μl of 0.85 % NaCl (w/v) in distilled water was added to each sample to make a final 1:10 dilution of serum. The inactivated serum was then diluted 1:2 across clear V-bottom plates (Sigma) to a final volume of 25 μl. Then, 4 HA units of virus diluted PBS to a final volume of 25 μl was added to each well. Lastly, 50 μl of 0.5% chicken RBC was added to each well. The plates were read after a 1-hour incubation at 4°C.

