## Supplementary Data for "Broadly-neutralizing antibodies that bind to the influenza hemagglutinin stalk domain enhance the effectiveness of neuraminidase inhibitors via Fc-mediated effector functions"

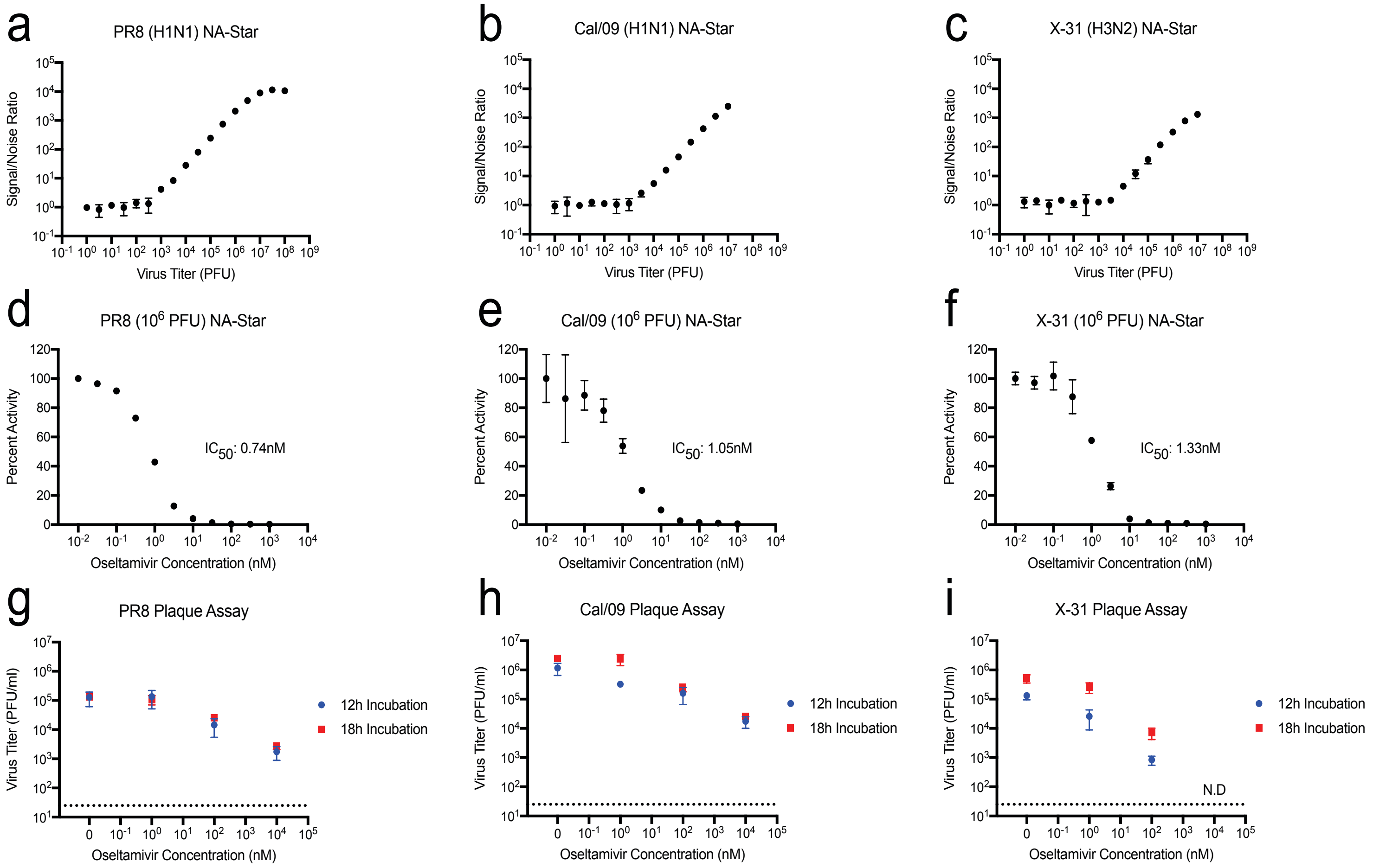


**Figure S1** Oseltamivir inhibits neuraminidase activity and replication of H1 and H3 influenza viruses.

(A-C) Neuraminidase activity of influenza viruses PR8, Cal/09, and X-31 were measured in-vitro using the NA-Star Neuraminidase Kit (ThermoFisher). Data shown as mean ± SD of at least two technical replicates.

(D-F) Oseltamivir susceptibility of the strains were then determined using $1\times10^6$ PFU of the three strains using the NA-Star Neuraminidase Kit (ThermoFisher). The $IC_{50}$ values are displayed to the right of the graphs. Data shown as mean ± SD of three technical replicates.

(G-I) Oseltamivir sensitivity of influenza virus strains was also quantified using plaque assays by infecting A549 cells with the virus at an MOI of 5 before incubating the infected cells with indicated concentrations of oseltamivir for 18 hours. N.D: not detectable. The limit of detection for the plaque assays was 25 PFU/ml, denoted by the horizontal dotted line. Data shown as mean ± SD of three technical replicates.

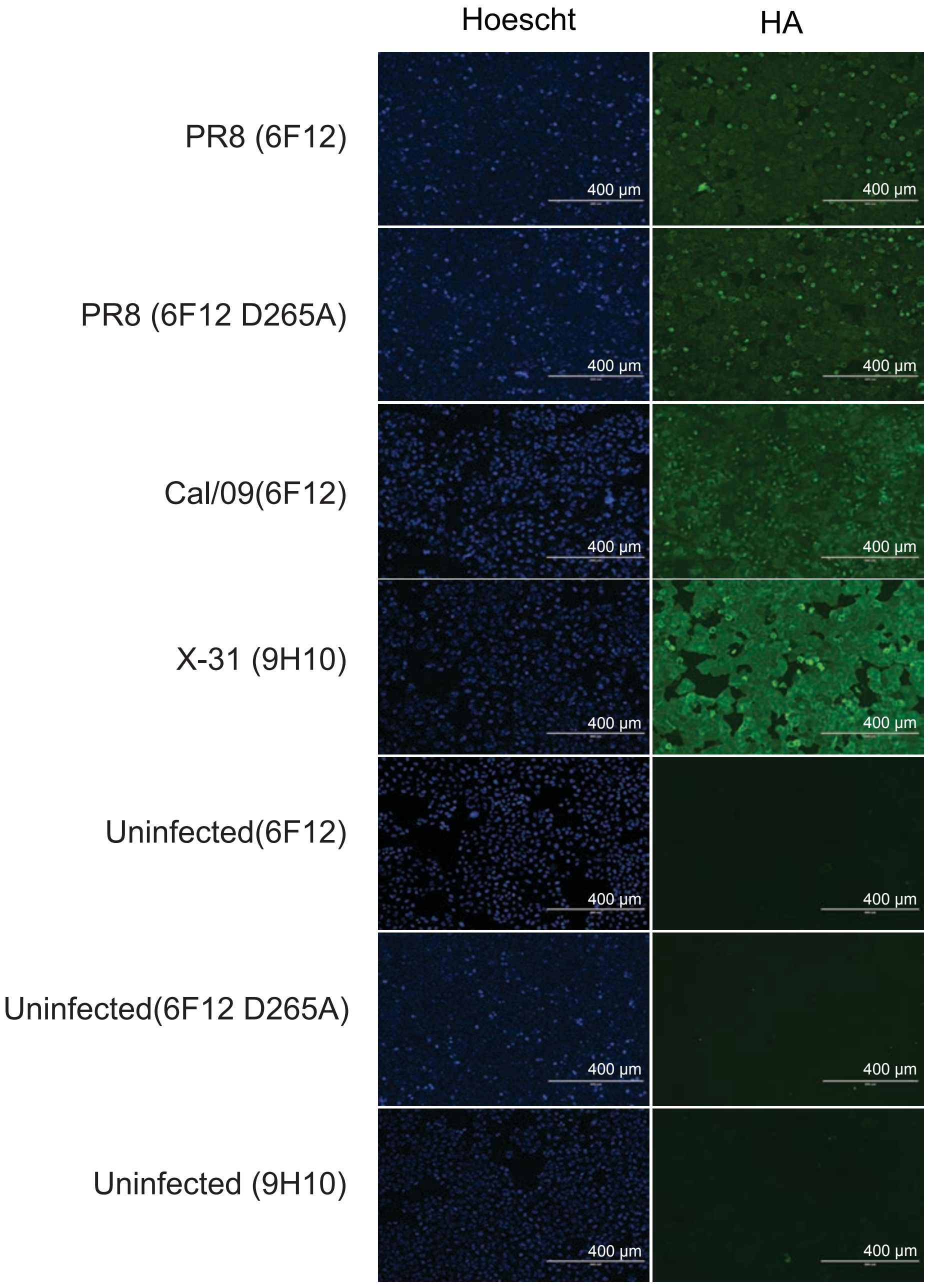


**Figure S2** Immunostaining of infected A549 cells by bNAbs.

A549 cells infected with PR8 or Cal/09 were stained with 6F12 or 6F12 D265A, and cells infected with X-31 were stained with 9H10. Uninfected A549 cells were stained using 6F12, 6F12 D265A, and 9H10 as negative controls. Scale bars are 400μm.

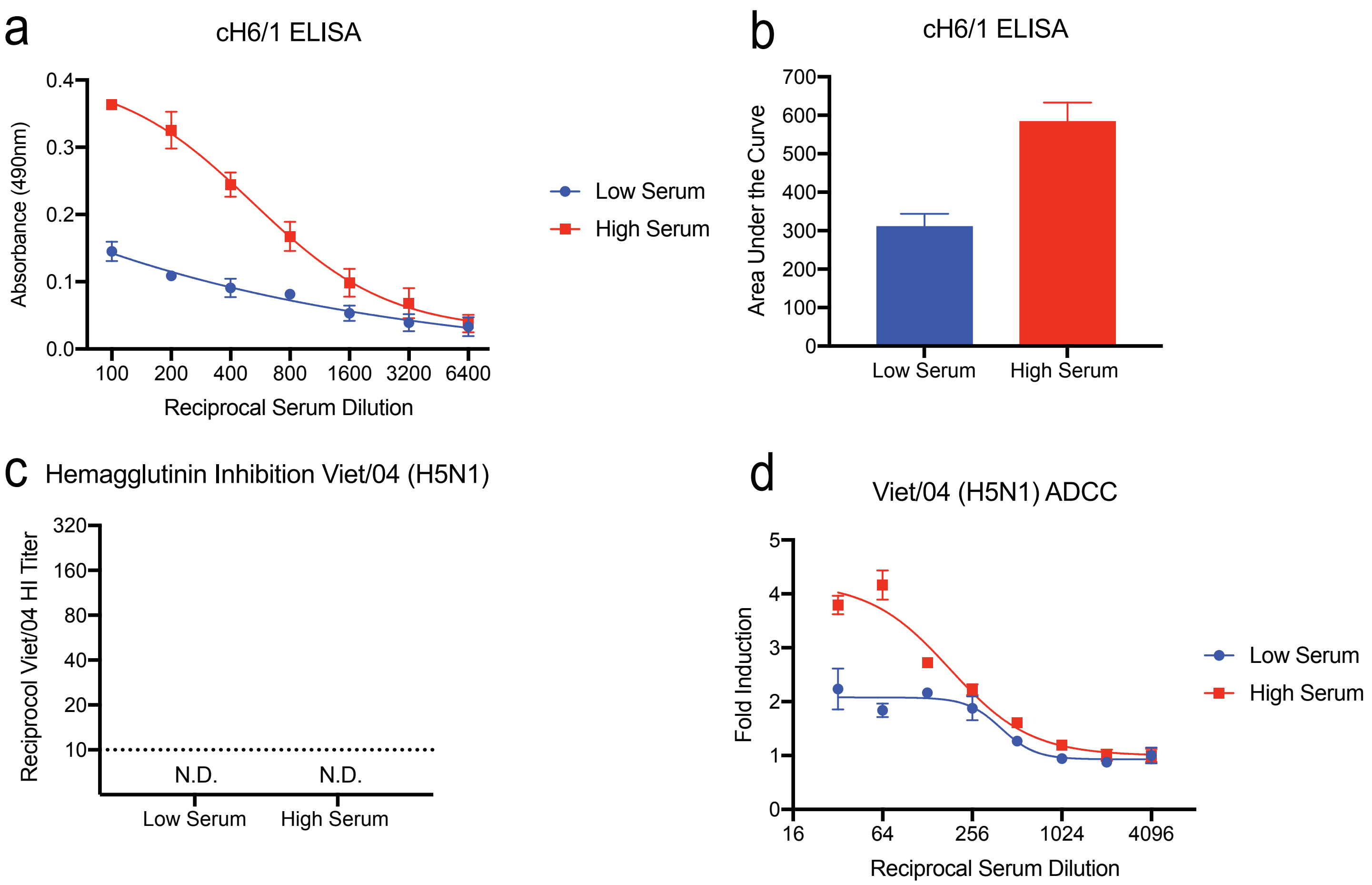


**Figure S3** Characterization of polyclonal stalk-binding antibodies in human serum. Human serum was obtained from peripheral blood of two healthy adult donors.

(A-B) ELISAs were performed using chimeric cH6/1 protein to quantify the titers of antibodies that bind to the stalk domain of H1 hemagglutinin. Absorbance data is shown as mean ± SD with technical triplicates. The area under the curve is also shown as mean ± SEM.

(C) Hemagglutinin inhibition assays using were used to test for the presence of antibodies inhibit receptor binding by the head domain of Viet/04 H5. N.D. = Not detectable.

(D) In vitro ADCC assays were completed using A549 cells infected with Viet/04 (H5N1) at an MOI of 5. Fold induction denotes activation above background (infected cells without antibody). Fold induction data is shown as mean ± SD with technical triplicates.
